# Development and validation of microsatellite markers from *de novo* transcriptome assembly of eggplant (*Solanum melongena* L.) and its putative progenitor *S. incanum* L. cultivars

**DOI:** 10.1101/560805

**Authors:** Shailesh K. Tiwari, Pallavi Mishra, Sakshi Singh, Vinay K Singh, Sarvesh P Kashyap, Major Singh, Kavindra N Tiwari, Prakash S Naik, Bijendra Singh

## Abstract

An elite cultivar of eggplant, Ramnagar Giant (*Solanum melongena* L.) and W-4 (*S. incanum* L.) with contrasting horticultural traits were used as parental lines to develop a mapping population of RILs. To accelerate breeding programs and to develop large scale SSR markers to be used in QTL mapping, RNA*Seq* libraries from different tissues of both the parental plants were deep sequenced and assembled into representation of a high quality *de novo* transcriptome using Illumina-based Next Generation Sequencing technology. 99.99% of high quality bases were obtained from all the tissues and deposited in TSA database at the NCBI link. Total 3, 156 and 3, 196 SNVs were detected in *S. melongena* and *S. incanum*, respectively. In *S. melongena*, 11, 262 SSR while in *S. incanum* 11, 829 SSR containing regions were identified. Based on functional annotation, 21, 914 unique genes could be identified for *S. melongena*, 21,706 unique genes for *S. incanum* and overall, 60 different transcription factors were identified in both the lines. Further, a total of 536 SSR markers were designed and screened for polymorphism of which, 157 markers produced polymorphism between the parental lines. The polymorphic SSRs shall be used for genotyping of RILs to map QTLs for various horticultural traits in eggplant and identification of candidate genes in response to biotic and abiotic stress.

## Introduction

Eggplant (*Solanum melongena* L., Solanaceae; *2n = 2x = 24*) commonly called brinjal or aubergine, is the third most important vegetable crop after tomato and potato in the genus *Solanum* [1]. While most of the other species of this genus are representatives of new world, eggplants are phylogenetically distinct and are indigenous to the old world with most related species reported to be originated in India, Africa, and South America [2, 3]. FAOSTAT data reveals that eggplant production has increased up to three folds in the last two decades, with China, India, Egypt, and Turkey being the top four eggplant producing countries across the world [4]. The berries are rich in naturally occurring water-soluble flavonoids [5], vitamins and minerals [6], polyphenol derivatives [7, 8], antioxidants like chlorogenic acid (CGAs) [9], caffeic acid [10, 11], and many other phytochemical compounds of human health benefits [12, 13]. Further, the low calorie, high potassium and high dietary fiber content makes it a vegetable of choice for the obese and diabetes patients [14].

High morphological diversity is observed among different cultivars and landraces of eggplant forms available in the market [15]. Common domesticated traits such as colour, shape, size, and chemical composition of fruits are highly specific to eggplant and important for breeding and marker-assisted selection (MAS) [16, 17]. The genetic variation in different morphological forms are a result of quantitative trait loci or QTLs which determine either complete or a portion of the total variation, and enables identification of large number of genes whose expression controls quantitative traits of agricultural importance such as plant yield, weight, fruit texture, quality, shape, color, chemical composition and plant resistance to diseases and pathogens [18]. Eggplant and their wild relatives have been an interesting subject of classical genetical analysis and research to the interested plant breeders [19]. Number of QTLs and mapping studies are reported for morphological traits in eggplant like fruit shape, size, colour, and prickliness [16, 17, 20], yield-related traits [21, 22], biochemical traits like location of CGA and polyphenol oxidase genes [9, 23], resistance to *Ralstonia solanaceaerum* [24], resistance to fungal wilts [25], resistance against Fusarium wilt [26], tolerance to bacterial wilt [27], and so on. Detailed biochemical and genetic information has been accumulated in eggplant for assessment of genetic diversity, QTL detection, and mapping based studies using range of molecular markers such as RAPD, AFLP, ISSR, SSR, SRAP, ESTs, and SNP [9, 16, 17, 26, 28–37]. These reports provide useful grounds for deciphering mechanisms underlying genetic control of the complex traits which are fundamental to the eggplant research program.

The progression of next-generation sequencing (NGS) technology has provided new ways for sequencing of plants especially for non-model species [38–40]. As compared to the traditional Sanger’s method for DNA sequencing, this method is cost-effective, fast, and useful in investigating the genome elements and molecular profile in comparatively easy ways [41]. NGS platform supports the view to sequence nucleotides frequently and allow massive sequencing of short sequence reads [42, 43] and large-scale development of molecular markers that can be helpful in analysing genetic diversity within large populations, validation of foreign gene introgression, linkage mapping, QTL mapping, *etc*. [41, 44]. Due to rapid advancements of whole-genome sequencing (WGS) and transcriptome methodologies in the last decade, assaying and gene expression profiling of many crops from the family *Solanaceae* were fastened to generate extensive molecular information. However, while the WGS of the model crops like tomato, potato, and pepper started early in 1990s and late 2000 [45–53], the eggplant genome has been merely addressed. Only a draft genome sequence of an eggplant cultivar, *Nakate-shinkuro* is published till date [54], which reflects that there is lack of a comprehensive dataset for eggplant genetic resources that could provide complete genetic information and makes the eggplant genome lagged behind with its natives with only little explored at genome and proteome level. The transcriptome assembly information in eggplant is also limited with only fewer reports published till date [1, 55–61]. The obtained information from published reports show that transcriptome data has wide applications in comprehensive characterization of entire genome, targeted sequencing of desired mRNA, discovery of novel transcription factor and identification of their binding sites, and expression profiling of non-coding mRNA sequences [44]. However, the genomic architecture and whole genome sequence information in eggplant is still lagging despite of extensive efforts to collect genetic and molecular information for this species.

Of the several species originated in Africa, *S. incanum*, a wild ally of eggplant is believed to be putative progenitor of *S. melongena* [3, 34–36], and together they both belongs to the complex *melongeneae*. They are readily crossable with each other to produce highly fertile interspecific hybrids and therefore are objects of interests in eggplant breeding and development of experimental populations. Therefore, with the objective of large-scale development of SSR markers for QTL mapping of domestication traits in eggplant and to collect more genomic information, we performed *de novo* transcriptome sequencing of two accessions of eggplant, Ramnagar Giant (*S. melongena* L.), a local cultivar restricted to Ramnagar and Varanasi regions of eastern Uttar Pradesh, and W-4, a genotype /line, a semi-wild close relative and its putative progenitor of eggplant *S. incanum* L., at the ICAR-IIVR Varanasi. The two accessions have been used as parents (Ramnagar Giant X W-4) to develop a mapping population of 114-F8 recombinant inbred lines (RILs). The parental lines are morphologically diverse from each other and exhibit distinct variations in traits of commercial and horticultural importance. W-4 is a spiny, wild form of eggplant showing resistance to some common pests hampering eggplant cultivation like *Fusarium oxysporum* f. sp. *Melongenae* [62, 63], *Phomopsis vexans* [64], and *Leucinodes orbonalis* [65]; while Ramnagar Giant is a local cultivar from Varanasi, the plants with dark green broad foliage bearing huge size rounded fruits of light green color with 3-4 fruits per plant and fruit weight 0.80-2.5 kg, and reported to possess moderate resistance to the *Phomopsis blight* of eggplant [66]. While the transcriptome assembly information of *S. incanum* has been previously reported [55], this is the first comprehensive report of *de novo* transcriptome sequencing of Ramnagar Giant which is an indigenous eggplant cultivar. Here, we successfully developed high quality SSR markers to be used in QTL mapping of horticultural traits and also described a set of differentially expressed annotated transcripts that shall further be used in discovery of novel gene families and transcription factors (TFs) associated with biosynthesis of active plant ingredients in response to biotic or abiotic stress and in defense related mechanisms in eggplant.

## Materials and methods

### Plant tissue collection and mRNA library preparation

*De novo* transcriptome sequencing of *S. melongena* and *S. incanum* (**Fig 1**) was performed in the present study. Both the lines have been maintained at the experimental farm of ICAR-Indian Institute of Vegetable Research, Varanasi. For deep mRNA sequencing, total RNA was extracted from following plant tissues each from *S. melongena* and *S. incanum*; root, stem, leaves, peduncle, floral buds, flower, calyx, and fruits. The tissues collected were immediately washed in 95% ethanol, hashed into small pieces, suspended in solution of 15 ml RNAlator (Ambion Cat#7020) and left for 3-4 h at 4°C. The samples were then transferred to 50 ml screw-capped falcon vials and stored at −80°C. The isolated RNA was quantified using Nanodrop spectrophotometer while other qualititative estimations were performed by Agilent High-sensitivity Bioanalyzer Chip. RNA Integrity Number (RIN) was calculated and finally, Illumina HiSeq-2000 platform was used to sequence the cDNA synthesised from high-quality RNA of both the genotypes.

**Fig 1.**
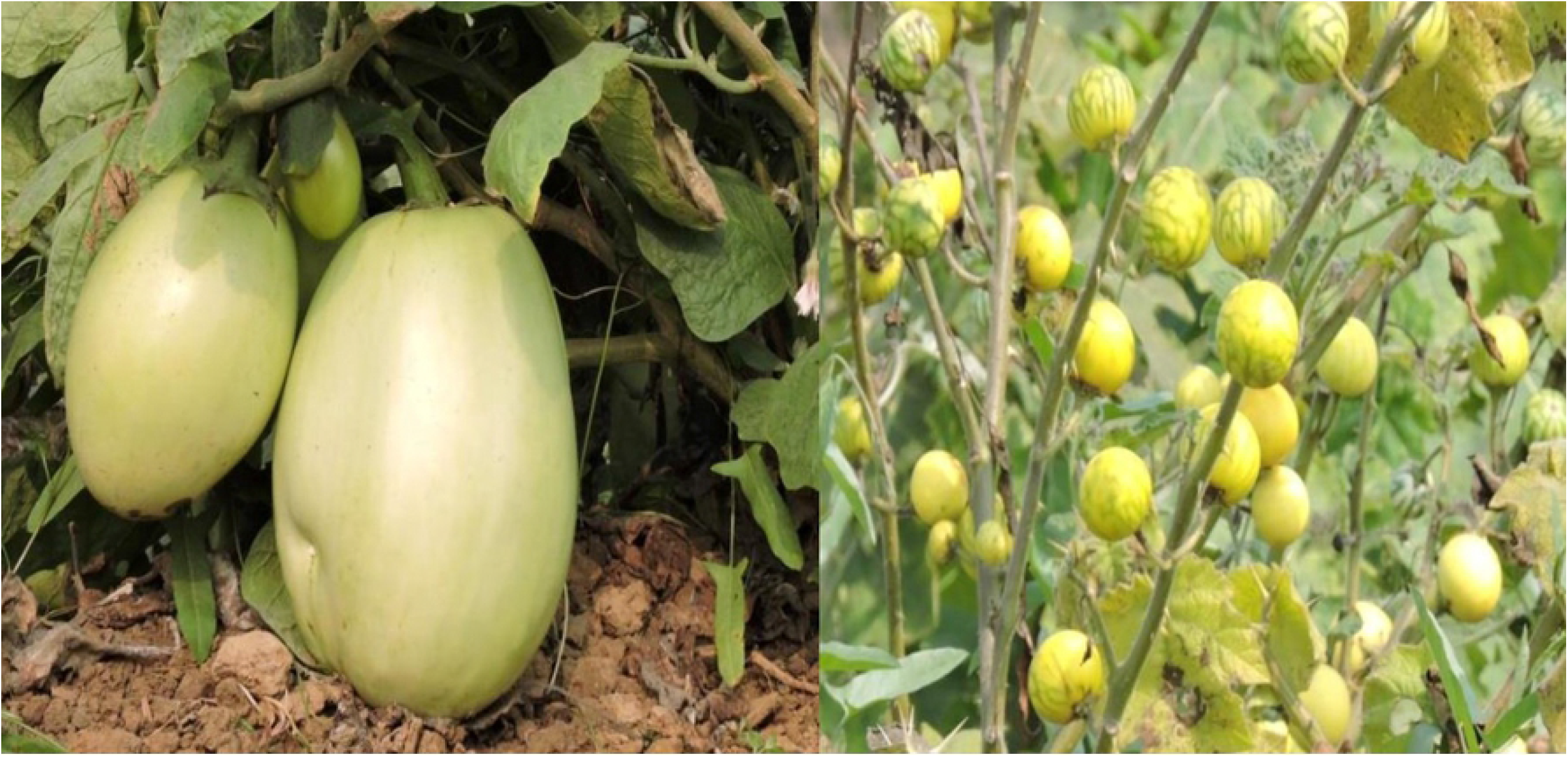
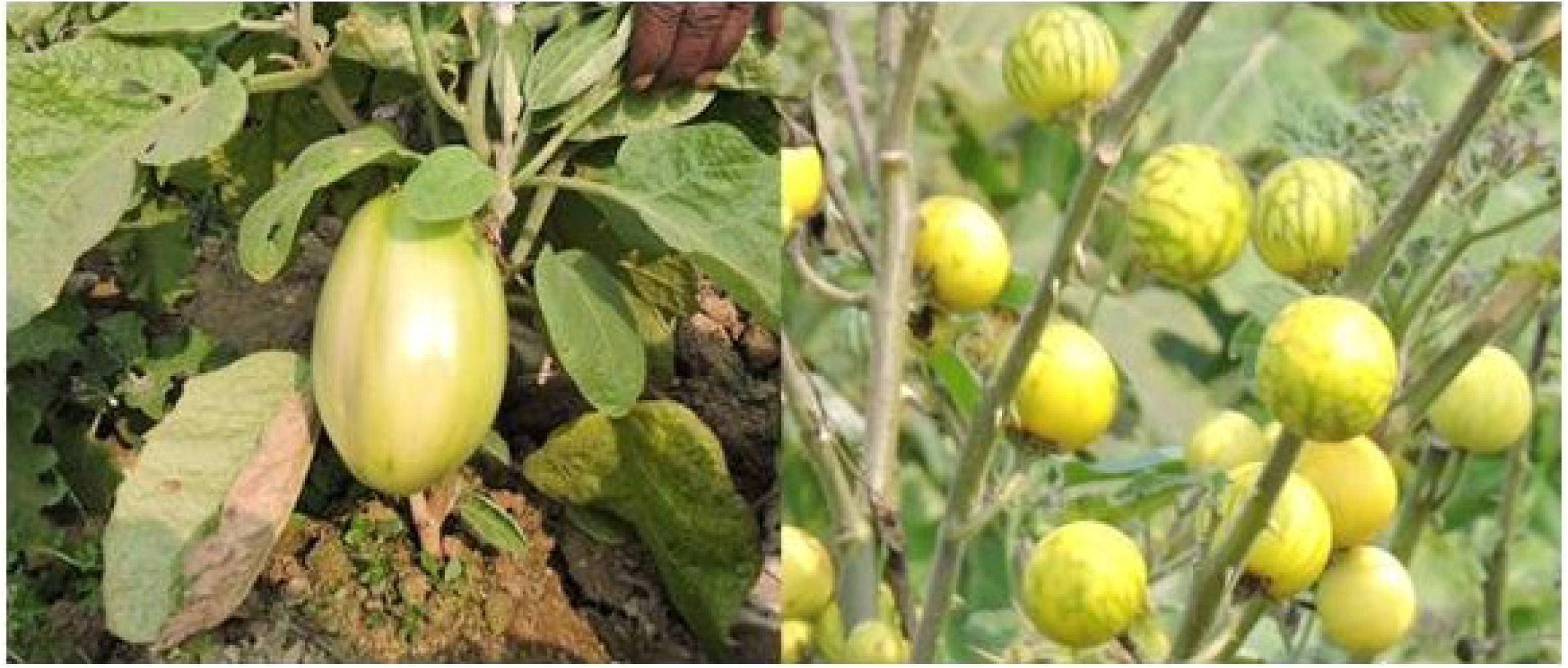
Ramnagar Giant (*S. melongena*) and W-4 (*S. incanum*) lines used for generating transcriptome assembly.

### Deep RNA sequencing and raw data quality control

Paired-end mRNA library was constructed from the total RNA as per the manufacturer’s protocol of Illumina based Next generation sequencer HiSeq-2000 to obtain high quality reads for their representation in the form of de novo assembly of the eggplant transcriptomes. Two cDNA libraries (read_1 and read_2) were constructed for the two genotypes on Illumina platform for transcriptome sequencing. More than 98% of high quality bases, both in forward and reverse (paired-end) reads were obtained from all the plant tissues. Based on this, the best quality reads (>200bp) were used for their assembly into contigs and further represented in the form of transcripts. All the sequencing related process such as quality testing of raw reads, filtering of High Quality (HQ) reads, trimming of adapter and low quality reads was done by SeqQC V2.0. The average Phred scaled quality score was found to be >20 which is an indication of a good quality of sequencing run.

### Transcriptome sequencing and NGS data assembly

Assembling of sequences through NGS technique allows one to obtain the high throughput fragments (reads) from the molecular (RNA) sequences which are further assembled and overlapped together to construct cDNA and can further be sequenced from both the directions to obtain paired-end reads [67]. The cDNA library from the pooled total RNA in equal quantity from all the plant tissues were sequenced from both the lanes to generate 100 bp paired-end reads with a coverage of 68.97x. *De novo* assembly of the obtained reads was accomplished with Velvet 1.2.10 and Oases v.0.2.08 using de Bruijn graph algorithm, and Oases (version 0.2.01) to obtain best quality raw data [68, 69]. The filtered transcripts (>200bp) were successfully deposited in Transcriptome Shotgun Assembly (TSA) database at the National Centre for Biotechnology Information (NCBI) portal with primary accessions GAYR00000000 and GAYS00000000 for *S. melongena* and *S. incanum*, respectively.

### SSR mining, SSR distribution, GC content, and SNV’s detection

SSR prediction was performed by the tool MISA v1.0 (MIcroSAtellite; http://pgrc.ipk-gatersleben.de/misa) [70], and analysis of GC content was carried out using SSR locator. Because SSR markers show greater degree of polymorphism and have co-dominant character, they are a good choice for diversity analysis and genetic mapping studies. The maximum fragment length considered for SSR prediction was 100 bp. For SNV’s detection and variation study, bowtie2 [71] and VarScan (v2.3.4) packages were used.

### Identification of TF classes

For identification of TF classes in *S. melongena* and *S. incanum*, sequence similarity search was done using PlantTFDB (plant transcription factor database), NCBI-CDD (conserved domain database) and tBLASTn algorithm taking *S. lycopersicum* TF sequences as the reference. For this, related protein sequences of a TF class was identified from Plant TFDB and aligned against the TSA of *S. melongena* (taxid: 4111) and *S. incanum* (taxid: 329799) using tBLASTn. Presence of conserved domains were further confirmed with NCBI-CDD (https://www.ncbi.nlm.nih.gov/Structure/cdd/)

### Similarity search, Functional annotation, and pathway analysis

Functional annotation of unigenes remains a difficult and challenging endeavour particularly for the non-model plant species such as eggplant due to limited stock of reference plant genomes at the public databases. For sequence similarity search and to predict annotation, the publically available databases BLASTx and TBLASTx search were performed to match the NCBI’s non-redundant (nr) plant sequences from the species including *S. lycopersicum, S. tuberosum, Vitis vinifera, Glycine max, Medicago truncatula* and *Ricinus communis* [72]. Homology search and motif search of the transcripts was executed using BLASTx. GO database and KAAS (KEGG Automatic Annotation Server: http://www.genome.jp/kegg/kaas/) server was used for gene annotation and pathways analysis, respectively [73, 74].

### Primer designing for polymorphism and QTL mapping in eggplant

MISA v1.0 was used for identification of SSR containing regions within 94, 838 sequences (spanning in length~73 Mb) in *S. melongena* and 95, 096 sequences (length~73 Mb) in *S. incanum*. A set of 536 SSR markers were designed using SSR locator and custom synthesized for detection of polymorphism between the parental lines (*S. melongena* and *S. incanum*), and further validation on the mapping population of RILs.

### DNA extraction and PCR protocol for newly synthesized SSR markers

Total genomic DNA was isolated from 20 days young leaves of *S. melongena* and *S. incanum* by CTAB method [75]. After isolation, 5 μl of DNA sample was mixed with 5 μl of 6X loading dye and run on an 0.8% agarose gel to access the DNA quality. DNA quantity was measured on Nanodrop spectrophotometer with peak absorbance at 260 nm. To check the amplification and validation of SSR primers upon parental lines and RILs, PCR reaction was carried out in a Biometra thermal cycler taking 50 ng genomic DNA as template mixed with 10 X Taq buffer, 2.5 mmol/l MgCl_2_, 0.2 mmol/l dNTPs, 0.5 mmol/l each of the SSR primer and 1 U Taq DNA Polymerase and final volume was made of 25 μl. The cyclic reactions involved 4 min at 94°C, 36 cycles of 1 min at 94°C, 1 min at 48°C, 2 min at 72°C, and finally 10 min at 72°C. The amplified products were separated on polyacrylamide gel electrophoresis (PAGE). For this, 8% non-denaturing gel was prepared in gel casters using 4 ml of 29:1 Acrylamide/Bis (% w/v), 8 ml distilled water, 1.68 gm Urea, 1.4 ml of 5 X TBE, 0.168 μl of 10% APS (ammonium persulphate) and 10 μl TEMED and left at room temperature for 45-60 minutes for polymerization to occur. The comb was slowly removed and gel was dipped in electrophoretic tank containing 10 X TBE buffer (pH 7.0-8.5) prepared at a working strength of 1 X. 10 μl PCR product (of both the parental lines and RILs) was loaded and separated for 3 h at 90 Volts. After separation, the gel was taken out carefully and stained with ethidium bromide in a separate tank and visualized under Alpha Imager ™ 3400 for genotyping of RILs and for further analysis.

## Results and discussion

### *De novo* transcriptome sequencing of *S. melongena* and *S. incanum*

After filtering adapter sequences and low quality sequences, a total of 101060300 numbers of raw reads were perpetuated from the raw data for sequencing, of which 50901025 number of HQ reads were obtained for *S. melongena* and 50159275 number of HQ reads were obtained for *S. incanum* (**Tables 1–3**). The raw paired-end read sequence data of FASTQ file size of **3.8 Gb** were obtained for *S. melongena* and *S. incanum*. On an average, the overall data quality was very good with more than 98% of HQ paired-end (PE) bases in both genotypes (**Figs 2A and 2B**). To assemble transcripts from short read sequences, Velvet and Oasis graph based assemblers were used [76]. 100.58 million (98.8%) HQ Paired reads were obtained for *S. melongena* and 100.32 million (98.86%) HQ paired reads were obtained for *S. incanum* (**Fig 3**). Bases having >= 20 phred score was considered for QC parameters in both the species. See raw and filtered data quality control report in **S1 Table**. The sequencing informations of *S. melongena* and *S. incanum* were submitted to the Sequence Read Archive (SRA) of NCBI portal with primary accessions SRX365148 and SRX365146, respectively.

**Fig 2A.**
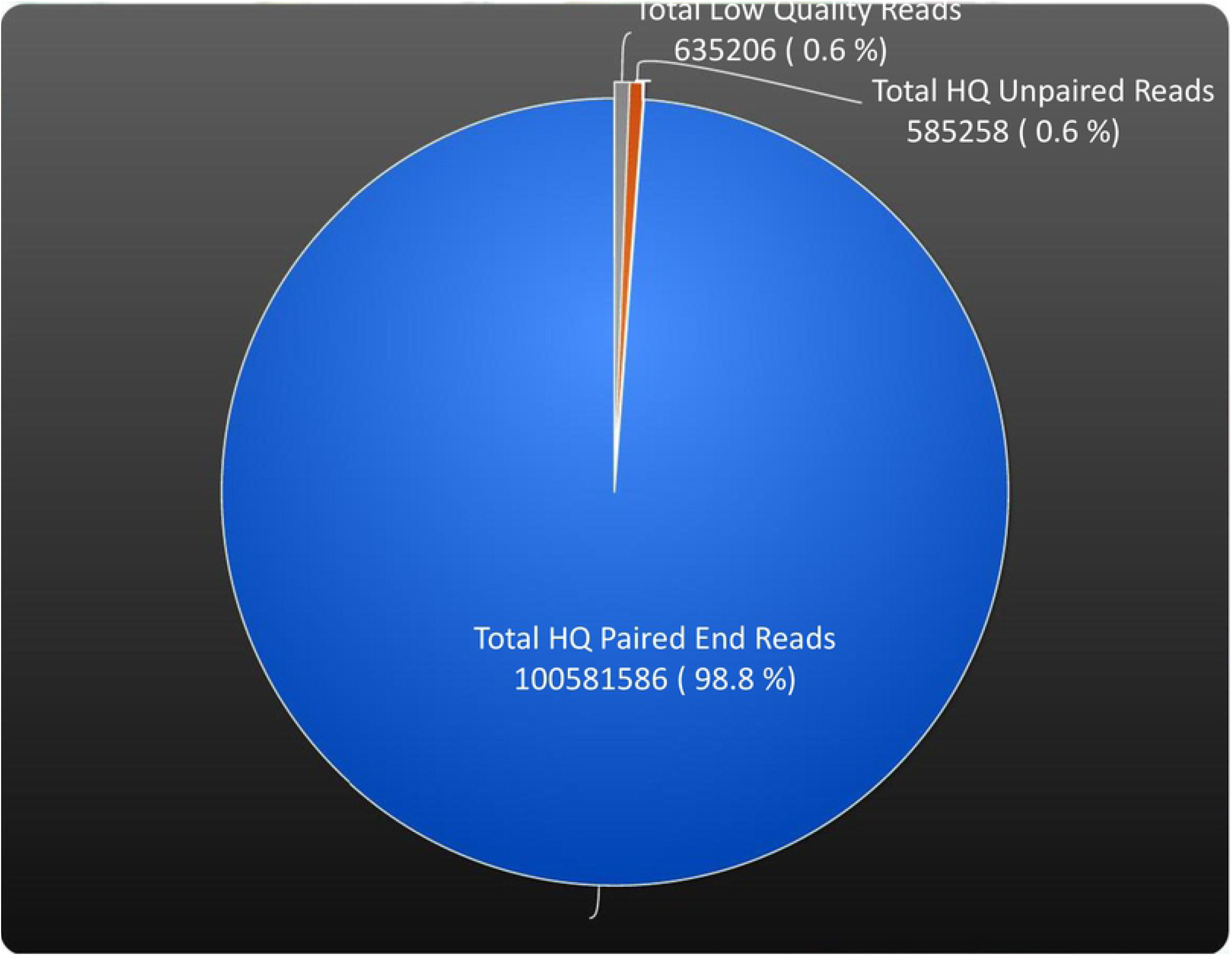
Paired end reads statistics of *S. melongena*.

**Fig 2B.**
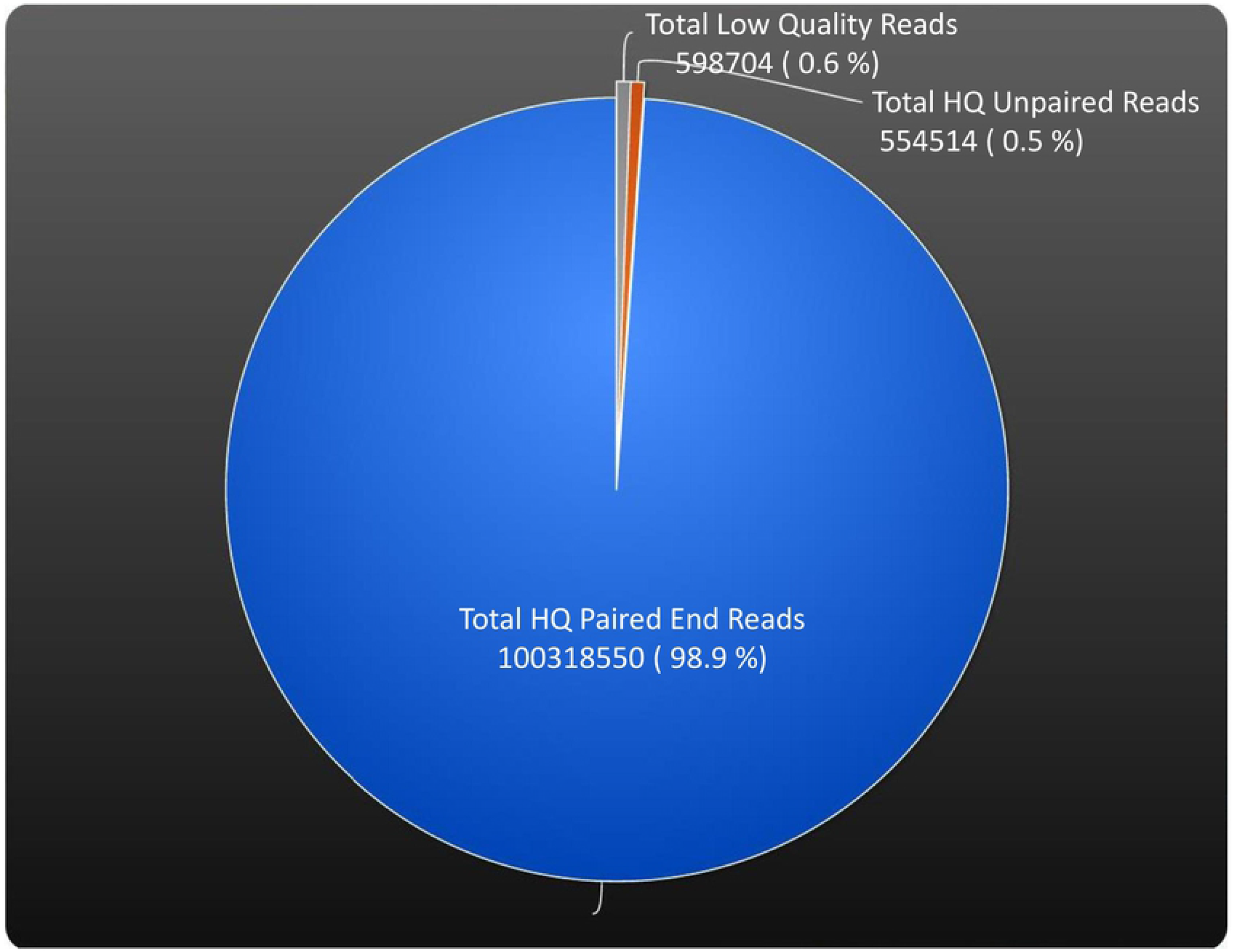
Paired end reads statistics of *S. incanum*.

**Fig 3.**
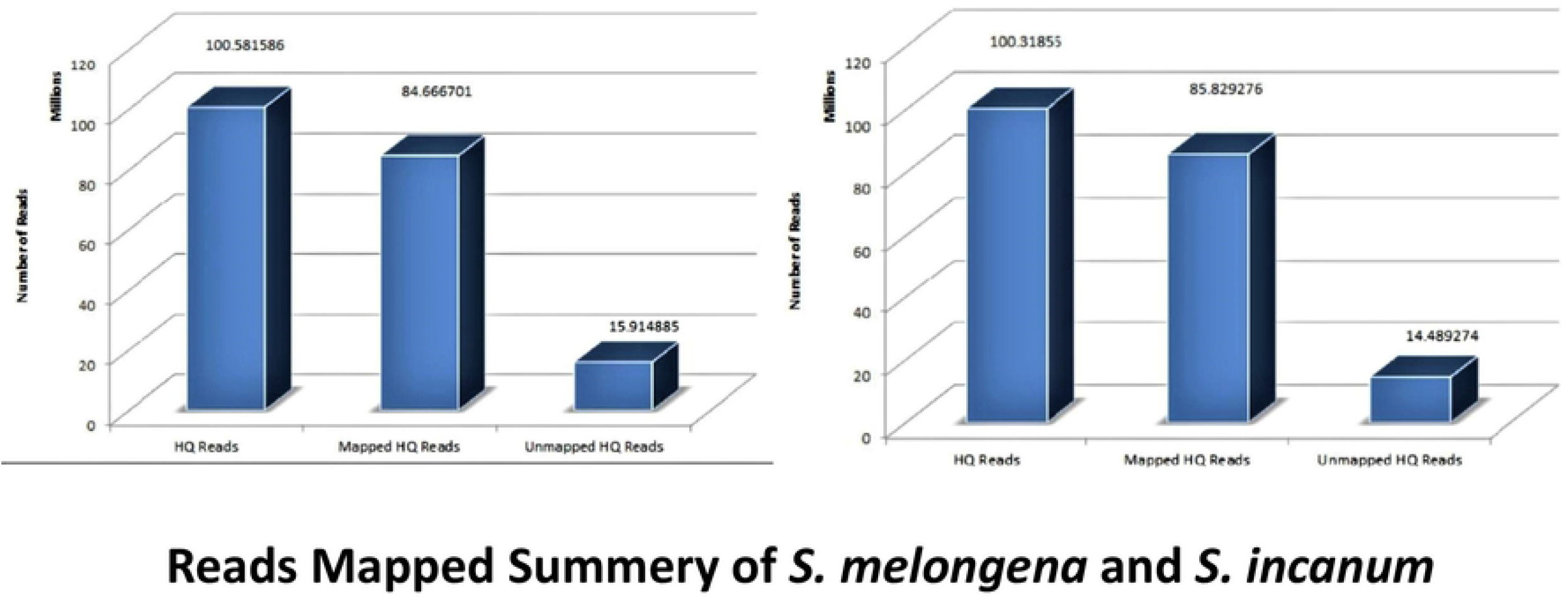
Graphical representation of total reads mapped summery.

**Table 1.** Assembly statistics of *S. melongena* and *S. incanum* transcriptomes.

| Parameters | *S. melongena* | *S. incanum* |
| --- | --- | --- |
| Total No. of Transcripts (>200bp) | 145512 | 144948 |
| Transcriptome Length(bp) | 149498096 | 147728785 |
| Min Transcript Length (bp) | 201 | 201 |
| Max Transcript Length (bp) | 11255 | 12393 |
| Average Transcript Length (bp) | 1027.4 | 1019.2 |
| N50 Contig Size (bp) | 1737 | 1715 |

**Table 2.**
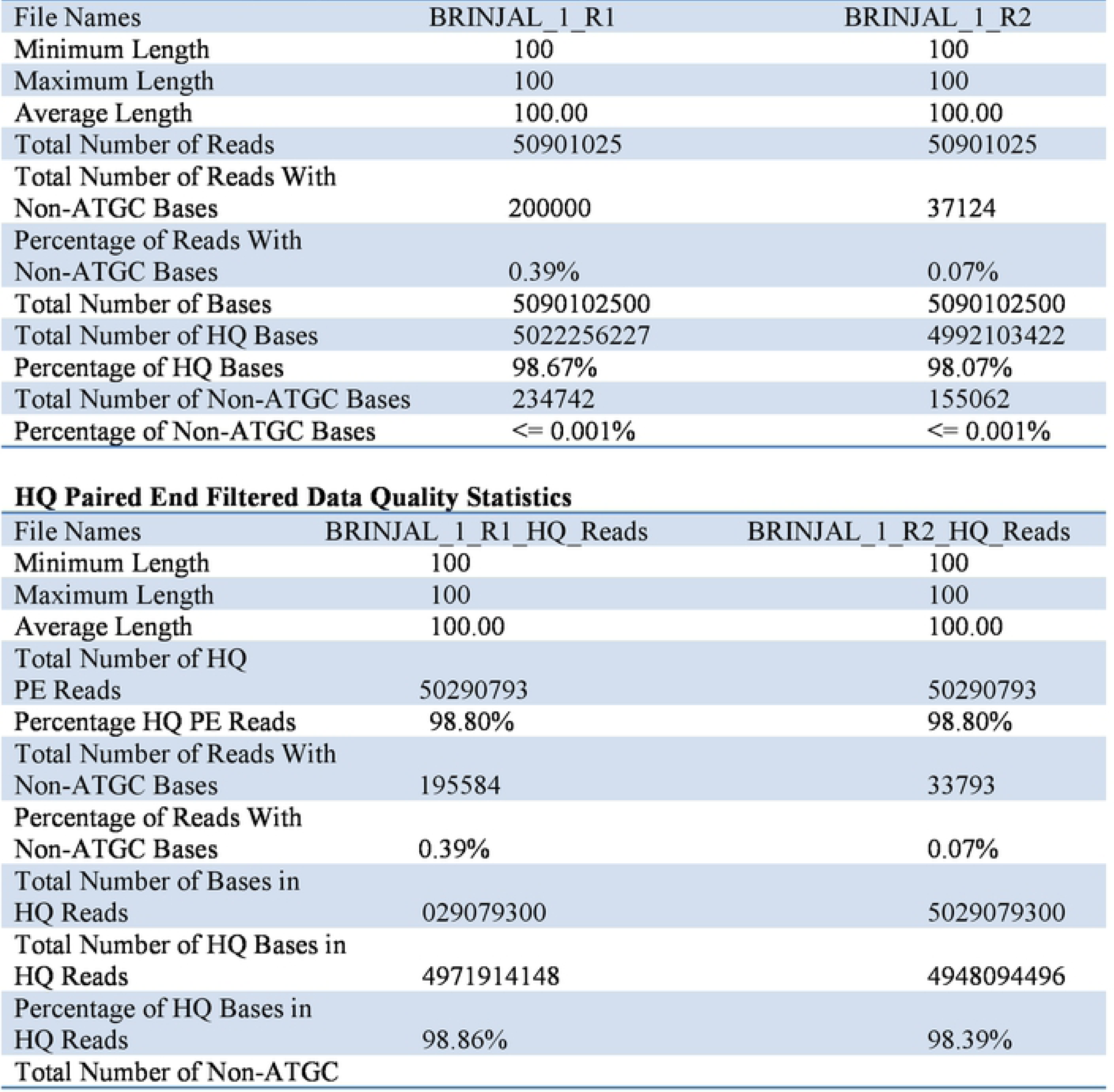
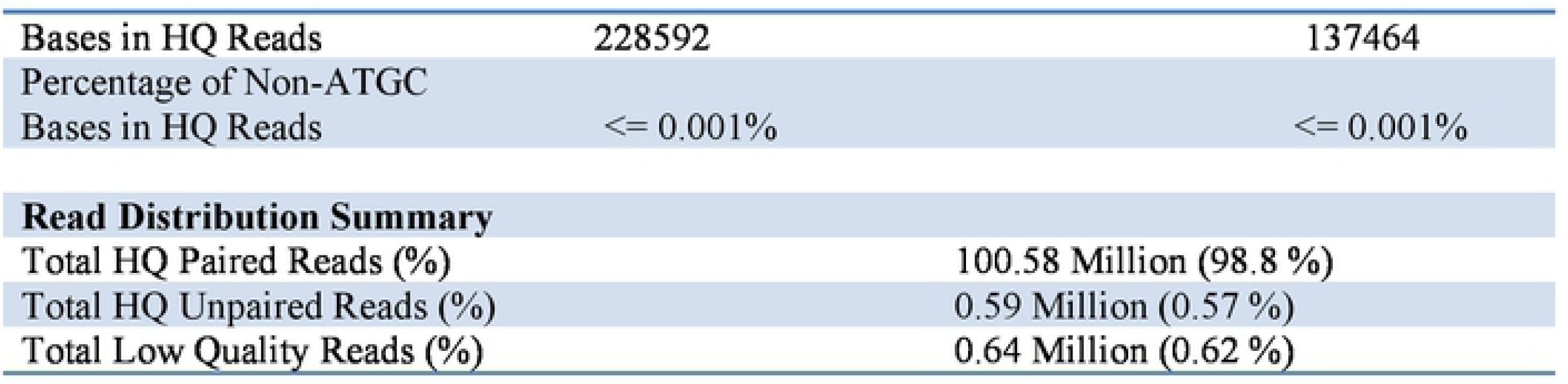
Raw Data Quality Statistics of *S. melongena*.

**Table 3.** Raw Data Quality Statistics of *S. incanum*.

| File Names | BRINJAL_2_R1 | BRINJAL_2_R2 |
| --- | --- | --- |
| Minimum Length | 100 | 100 |
| Maximum Length | 100 | 100 |
| Average Length | 100.00 | 100.00 |
| Total Number of Reads | 50735884 | 50735884 |
| Total Number of Reads With Non-ATGC Bases | 165813 | 38016 |
| Percentage of Reads With Non-ATGC Bases | 0.33% | 0.07% |
| Total Number of Bases | 5073588400 | 5073588400 |
| Total Number of HQ Bases | 5009410203 | 4981220984 |
| Percentage of HQ Bases | 98.74% | 98.18% |
| Total Number of Non-ATGC Bases | 178048 | 158746 |
| Percentage of Non-ATGC Bases | <= 0.001% | <= 0.001% |

**HQ Paired End Filtered Data Quality Statistics**
| File Names | BRINJAL_2_R1_HQ_Reads | BRINJAL_2_R2_HQ_Reads |
| --- | --- | --- |
| Minimum Length | 100 | 100 |
| Maximum Length | 100 | 100 |
| Average Length | 100.00 | 100.00 |
| Total Number of HQ PE Reads | 50159275 | 50159275 |
| Percentage HQ PE Reads | 98.86% | 98.86% |
| Total Number of Reads With Non-ATGC Bases | 162235 | 34852 |
| Percentage of Reads With Non-ATGC Bases | 0.32% | 0.07% |
| Total Number of Bases in HQ Reads | 5015927500 | 5015927500 |
| Total Number of HQ Bases in HQ Reads | 4961753705 | 4939544661 |
| Percentage of HQ Bases in HQ Reads | 98.92% | 98.48% |
| Total Number of Non-ATGC Bases in HQ Reads | 173950 | 141882 |
| Percentage of Non-ATGC Bases in HQ Reads | <= 0.001% | <= 0.001% |

| Read Distribution Summary |  |
| --- | --- |
| Total HQ Paired Reads (%) | 100.32 Million (98.86 %) |
| Total HQ Unpaired Reads (%) | 0.55 Million (0.55 %) |
| Total Low Quality Reads (%) | 0.6 Million (0.59 %) |

### SSR identification in *S. melongena* and *S. incanum*

Using the SSR locator and MISA v1.0 tool, SSR containing regions were identified within 94,838 sequences (spanning in length~73 Mb) in *S. melongena* and 95,096 sequences (spanning in length~73 Mb) in *S. incanum*. Maximum number of bases interrupting two consecutive SSRs in a compound microsatellite was found to be 100 bp. Total numbers of 11,262 and 11,829 SSRs could be identified in *S. melongena* and *S. incanum*, respectively. These SSRs were distributed in 9,839 numbers of sequences in *S. melongena* and 10,235 sequences in *S. incanum*. In *S. melongena*, highest percentage of SSR distribution was accounted for mononucleotides (~58%), followed by trinucleotides (~22%), dinucleotides (~15%) and complex, tetra- and penta-nucleotides approximately 4%, 1% and 0.12%, respectively and GC content was found to be 43%. In *S. incanum*, mononucleotides accounted for ~54%, followed by trinucleotides (~21%), dinucleotides (~14%) and complex, tetra- and penta-nucleotides approximately 5%, 3% and 0.8%, respectively and 43% GC content (**Fig 4 and Tables 4 and 5**). The distribution of SSRs and their motifs are summerized in **S2 and S3 Tables**. Through SSR locator, a total of 536 new SSR markers were designed and custom synthesized from the transcriptome sequence of *S. melongena* and *S. incanum*. (Details and sequences of all primers are given in **S2 Table**).

**Fig 4.**
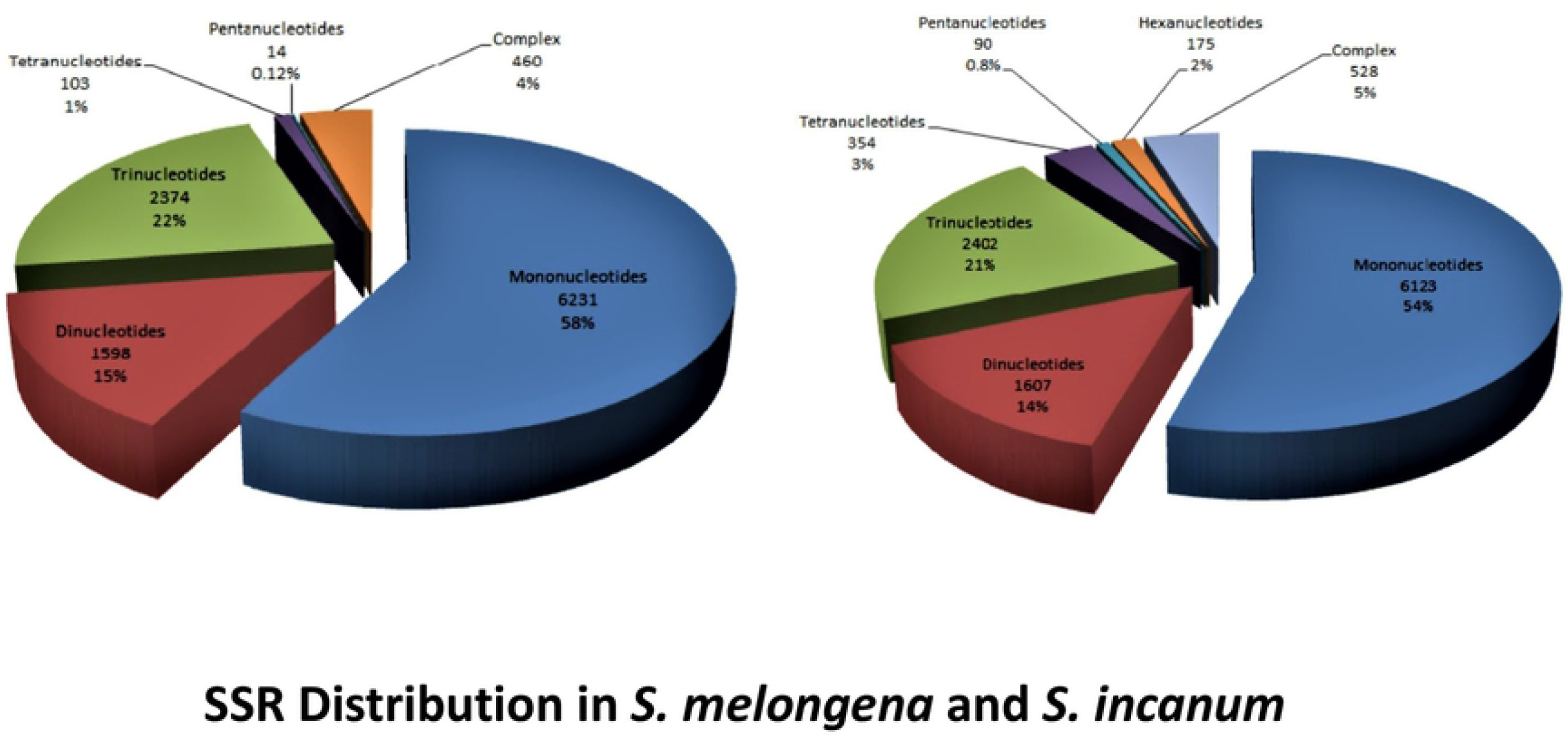
Graph showing identified SSR with distribution.

**Fig 5.**
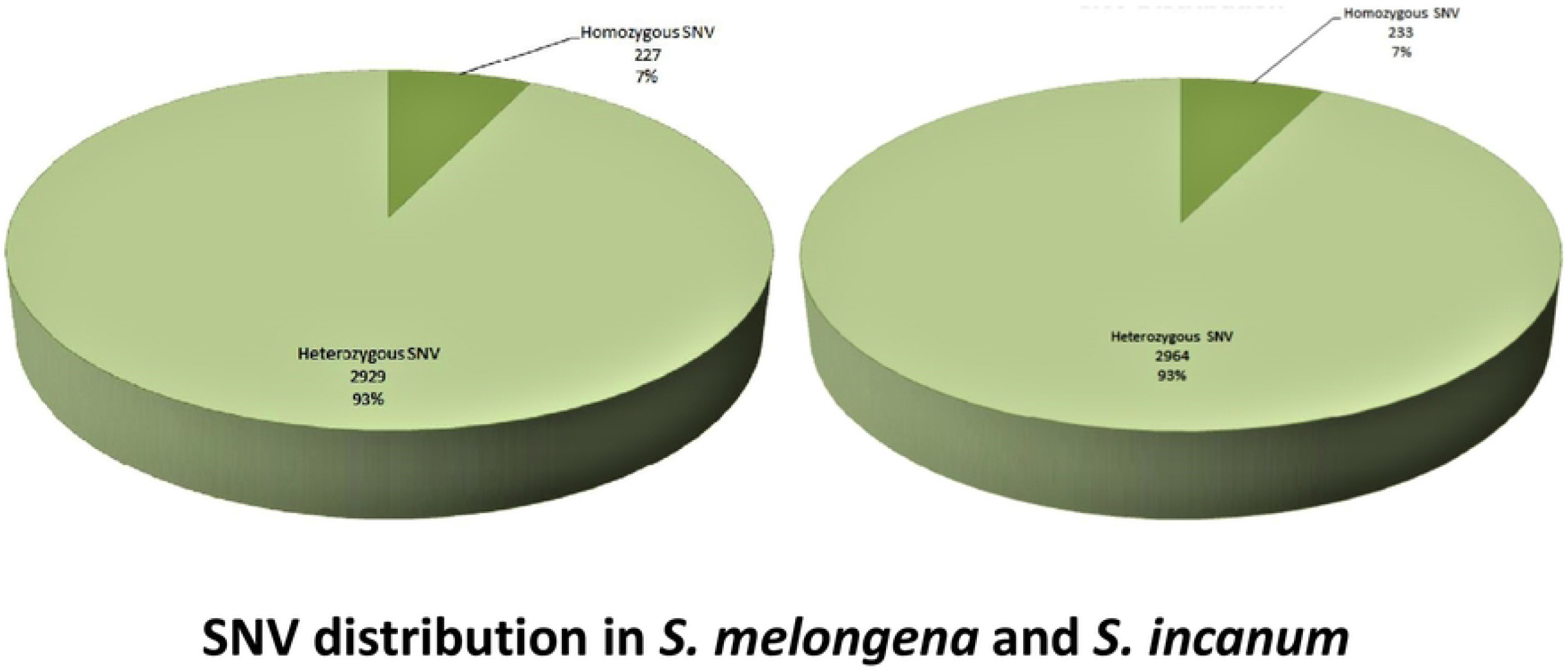
Graph showing SNV distribution.

**Table 4.** Statistics of SSR identified in *S. melongena* and *S. incanum*.

| Unit size of microsatellite | Minimum number of repeats |
| --- | --- |
| Mono-nucleotide repeats | 10 |
| Di-nucleotide repeats | 6 |
| tri-, tetra-, penta- & hexa-nucleotide repeats | 5,4,4,4 |
| Maximal number of bases interrupting 2 SSRs in a compound microsatellite | 100 |

| Statistics of microsatellite search for *S. melongena* and *S. incanum* |  |  |
| --- | --- | --- |
| Parameters / SSR motif length | *S. melongena* | *S. incanum* |
| Total number of sequences examined | 94,838 | 95096 |
| Total size of examined sequences (bp) | 73634990 (~73MB) | 73796620 (~73MB) |
| Total number of identified SSRs | 11262 | 11829 |
| Number of SSR containing sequences | 9839 | 10235 |

**Table 5.** Distribution of different SSR type classes in *S. melongena* and *S. incanum*.

| SSR class | Number of SSRs Detected |  | % of SSRs Detected |  |
| --- | --- | --- | --- | --- |
|  | *S. melongena* | *S. incanum* | *S. melongena* | *S. incanum* |
| Mononucleotides | 6231 | 6123 | 57.80 | 54.62 |
| Dinucleotides | 1598 | 1607 | 14.82 | 14.33 |
| Trinucleotides | 2374 | 2402 | 22.02 | 21.43 |
| Tetranucleotides | 103 | 354 | 0.96 | 3.16 |
| Pentanucleotides | 14 | 90 | 0.13 | 0.80 |
| Hexanucleotides | - | 175 | - | 1.56 |
| Complex | 460 | 528 | 4.27 | 4.10 |

### SNVs detection

Bowtie2 tool was used for SNV’s detection and VarScan package (v2.3.4) was used for variation analysis to identify allelic variations in *S. melongena* and *S. incanum*. We reported a total number of 3, 156 SNVs in *S. melongena* and 3, 196 SNVs in *S. incanum*. Within these SNVs, 227 homozygous SNVs (~7%) were detected in *S. melongena* and 233 homozygous SNVs (~7%) were detected in *S. incanum*. The number of heterozygous SNVs detected were 2, 929 (~93%) and 2, 964 (~93%) in *S. melongena* and *S. incanum*, respectively (**Fig 5**). SNVs were further distinguished into separate classes based on within sequence variations (**Fig 6**). For other details, see **S4 Table**.

**Fig 6.**
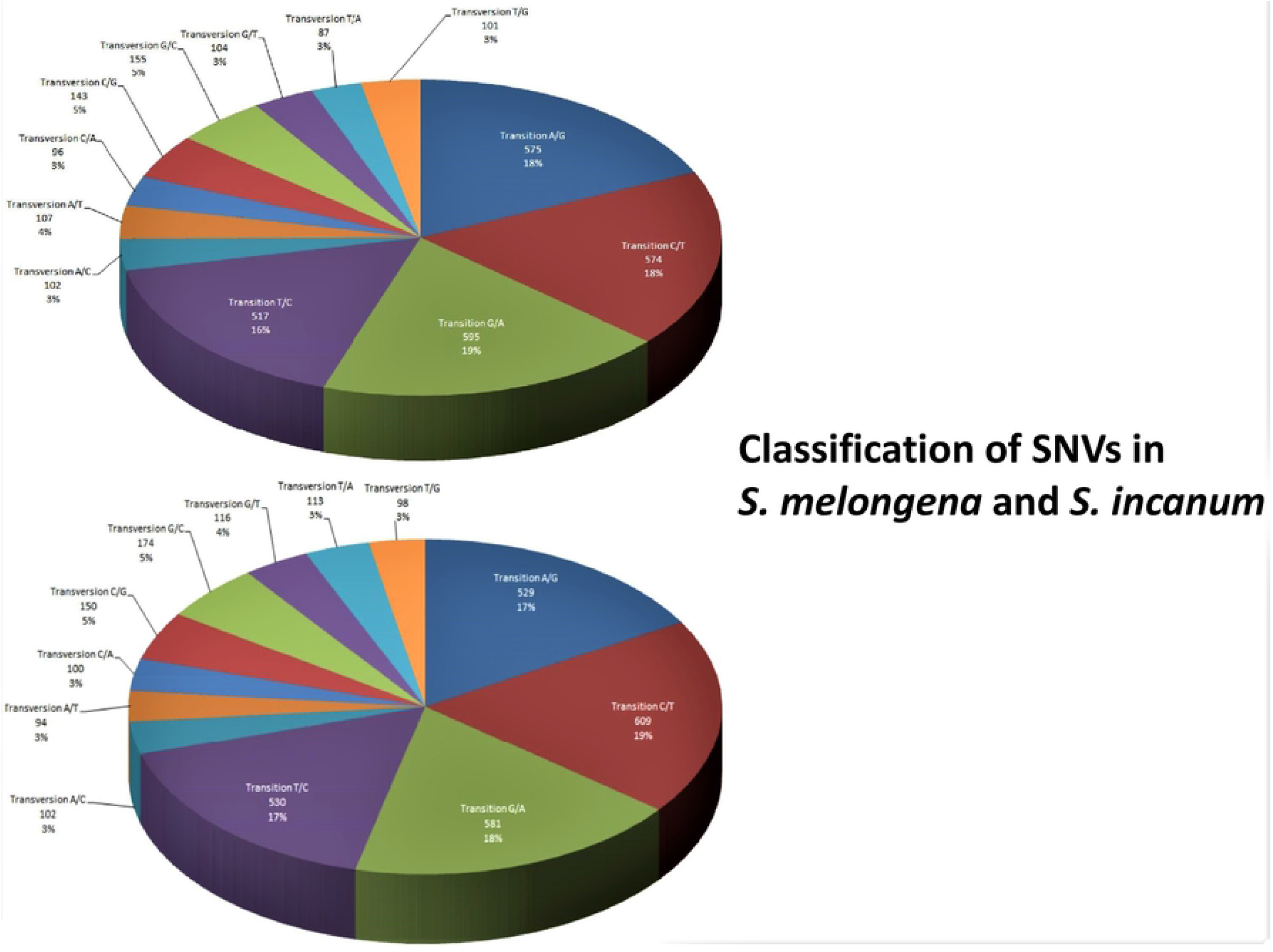
Graphical representation of variation classification.

### Transcripts annotation of *S. melongena* and *S. incanum*

Nucleotide non-redundant data sets from plant species like *S. lycopersicum, S. tuberosum, Vitis vinifera, Glycine max, Medicago truncatula* and *Ricinus communis* were used to annotate the clusterd transcripts to represent in the form of a library using BLASTx. A transcript coverage of cut off > 70% (E-value<= 0.001) was considered for homology prediction and functional annotation of eggplant transcripts. In *S. melongena*, total 94,838 predicted transcripts were examined to match against the Plant GDB for annotation of which 26, 914 transcripts (28.37%) could be annotated. Similarly, in *S. incanum*, 95, 096 predicted transcripts were examined to match against the Plant GDB of which 26, 660 transcripts (28.03%) could be annotated (**Fig 7**). Besides these, 21, 914 unique genes for *S. melongena* and 21, 706 unique genes for *S. incanum* were also identified (**S6 and S7 Tables**).

**Fig 7.**
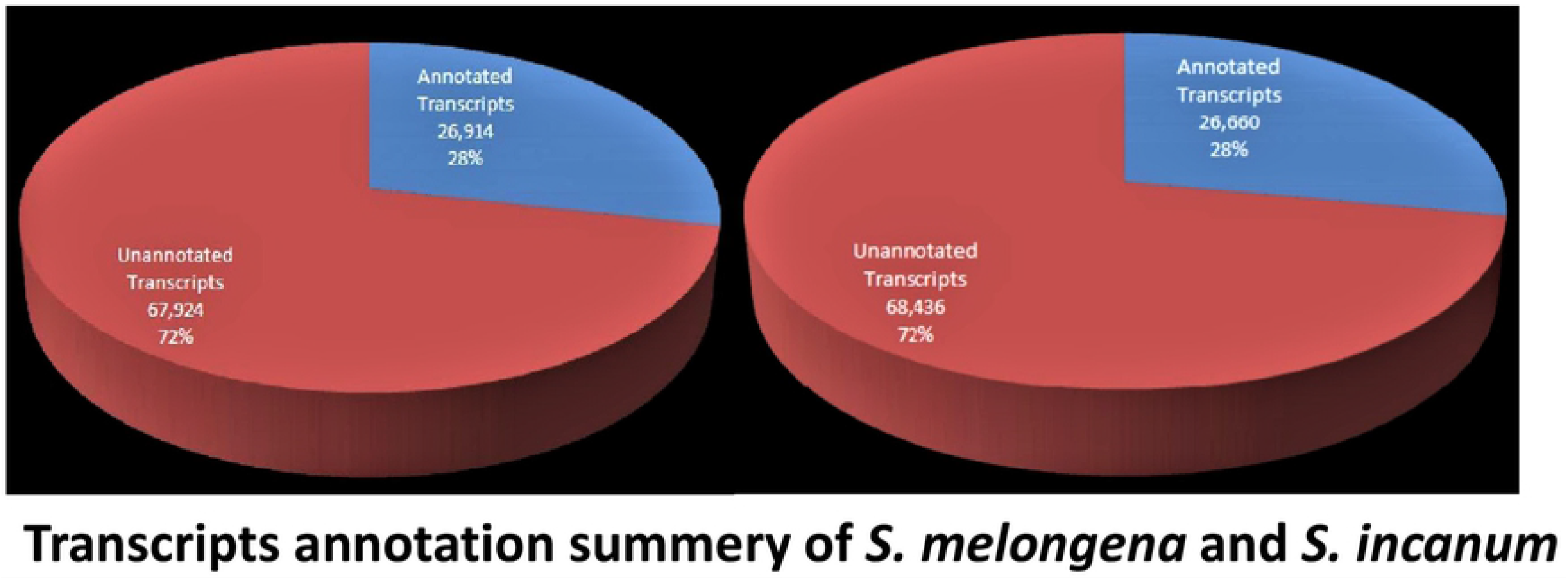
Transcript annotation summery of *S. melongena* and *S. incanum*.

### DEGs, Gene ontology, Unigene identification and pathway analysis

It was found that 379 genes were up-regulated, 368 genes were down-regulated, while 747 genes were found to be differentially expressed between the transcriptome assembly of *S. melongena* and *S. incanum*, respectively (**Fig 8**). A total of 21, 914 and 21, 706 unique genes were mapped in *S. melongena* and *S. incanum*, respectively. Analysis for identification of transcripts involved in cellular and biochemical pathways was carried out was based on KAAS (KEGG Automatic Annotation Server: http://www.genome.jp/kegg/kaas/) server. GO database was exploited for gene annotation in which, predicted transcripts with gene ontology annotation was 5, 087 out of 9, 085 annotated transcripts in *S. melongena and* 5, 064 out of 8, 968 annotated transcripts in *S. incanum*. The GO annotation summery of biological processes, cellular components, and molecular functions of *S. melongena* and *S. incanum* are described separately (**Fig 9 and S9 Table**). It was found that 1, 211 (out of total 9,085 annotated transcripts) in *S. melongena* and 937 (out of 8,938 annotated transcripts) in *S. incanum* could be assigned with KEGG pathway. The top ten KEGG pathways were identified in both the species based on KEEG ortholog, which play a key role in biochemical and cellular metabolism (**S8 and S9 Tables**).

**Fig 8.**
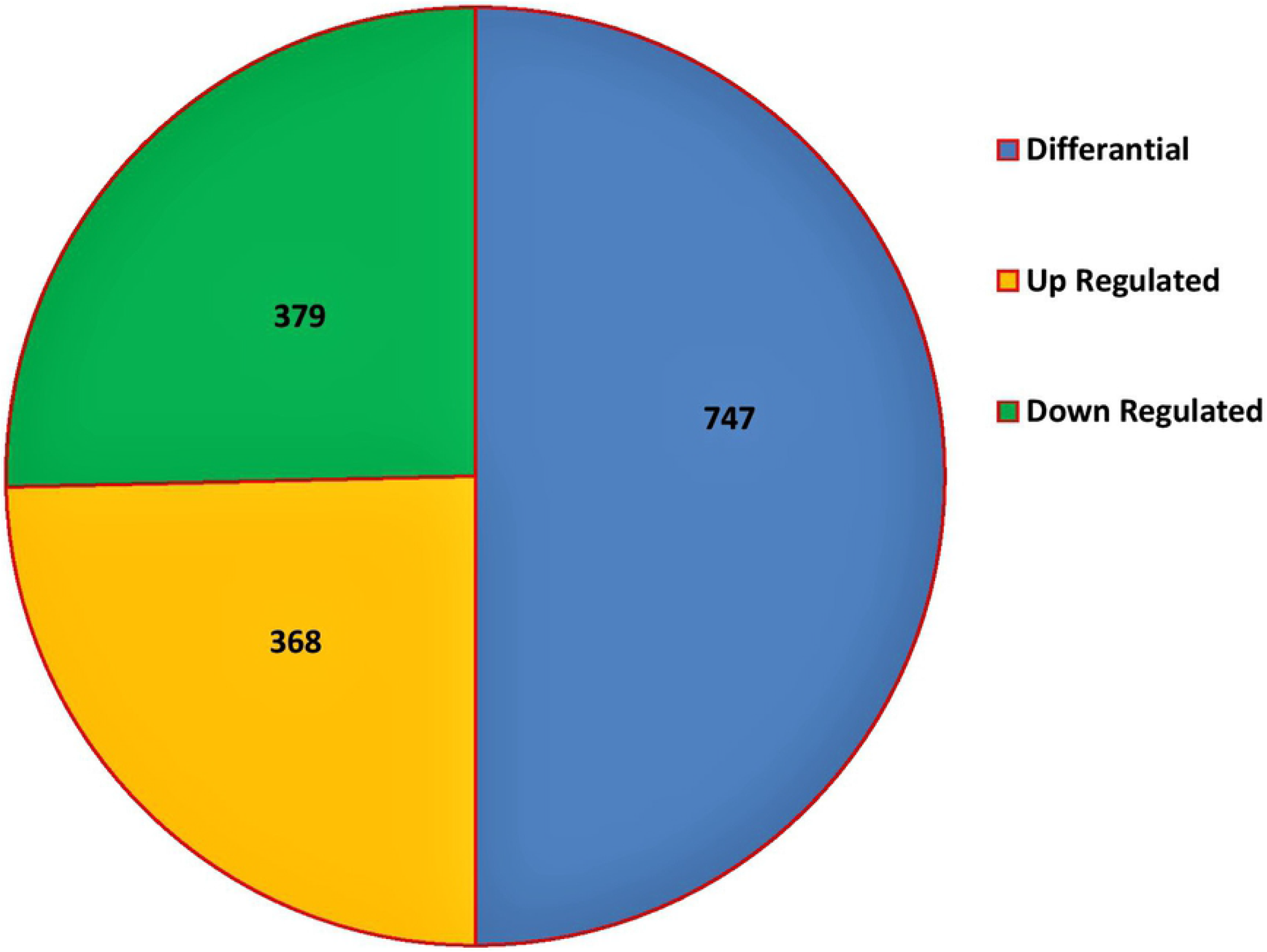
Differentially expressed genes details of transcripts.

**Fig 9.**
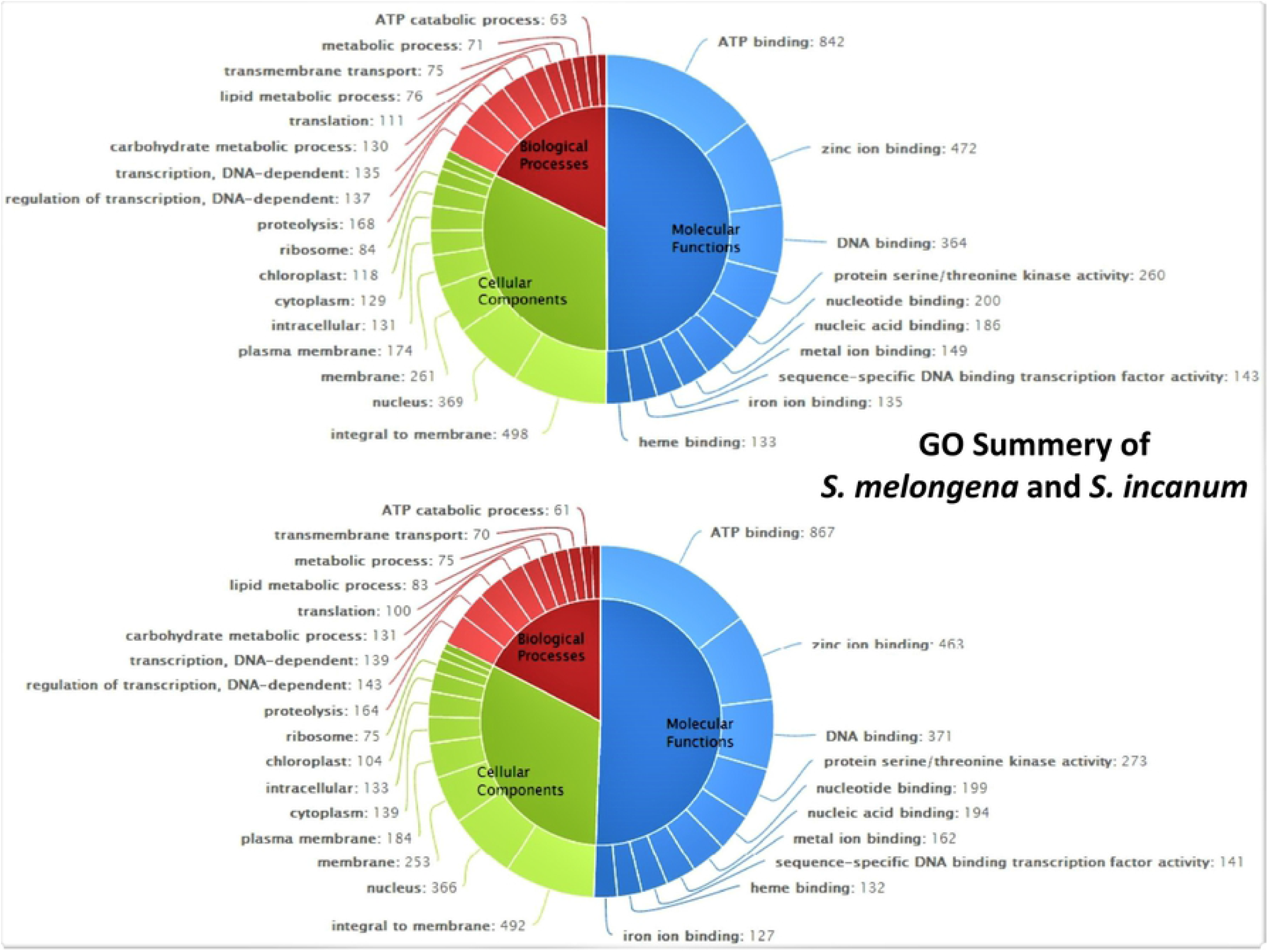
GO summery of *S. melongena* and *S. incanum*.

### Identification of TF classes in *S. melongena* and *S. incanum*

Overall, 60 different TF classes were identified in responses to biotic and abiotic stresses in transcriptome sequences of *S. melongena* and *S. incanum* which play a crucial role in different expression and developmental pathways. Each TF class represented number of genes identical to those available in PlantTFDB. Based on similarity search, some of the important TF associated genes such as *AP2, ARF, ARR-B, bHLH, bZIP, CAMTA, C2C2, Dof, ERF, GATA, WRKY, NAC* and *YABBY* were successfully identified in both the species (**Fig 10 and S10 Table**).

**Fig 10.**
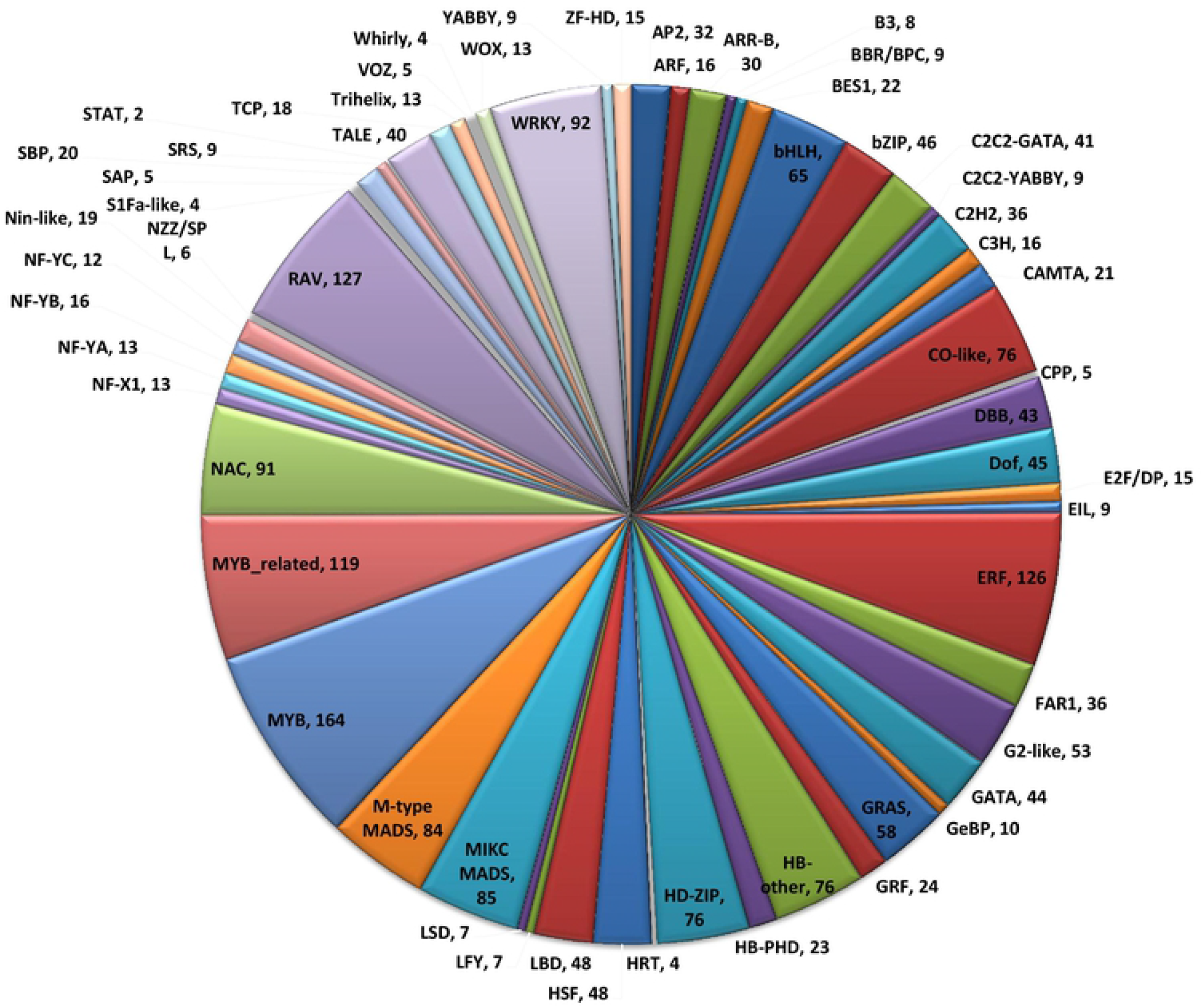
Identified transcription factor detail statistics.

### Parental screening for presence of polymorphism and validation of polymorphic SSRs on RILs

536 newly synthesized SSR primers were tested on parental lines to check the amplification of which 502 primers (93.65%) produced amplicons and 157 markers (31.27%) detected polymorphism between the parental lines through PAGE. The polymorphic SSRs were further validated on RILs for genotyping through PAGE (**Figs 11A and 11B**). The details of primers pairs used for detection of polymorphism between parental lines and obtained polymorphic primers are given in **S2 Table**, and have been used with suffix ‘*Sm*’ and ‘*Si*’ for primer sequence IDs of *S. melongena* and *S. incanum*, respectively for further studies. From the obtained results, it can be inferred that NGS-derived markers are more useful and informative in detecting genetic polymorphism between related and highly inbred species. Further, the NGS data can be exploited in much broader sense to collect more genomics and proteomics information for non-model species like eggplants.

**Fig 11A.**
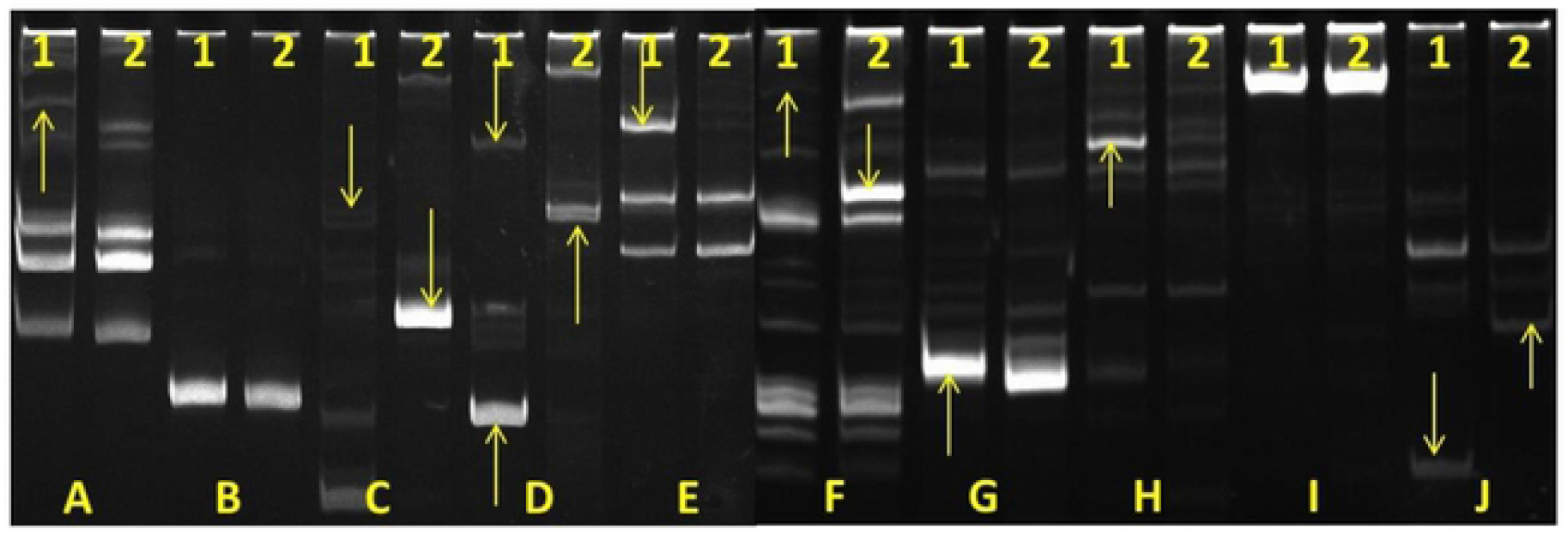
PAGE images showing different transcriptome derived SSR markers used for screening polymorphism between parental lines.

**Fig 11B.**
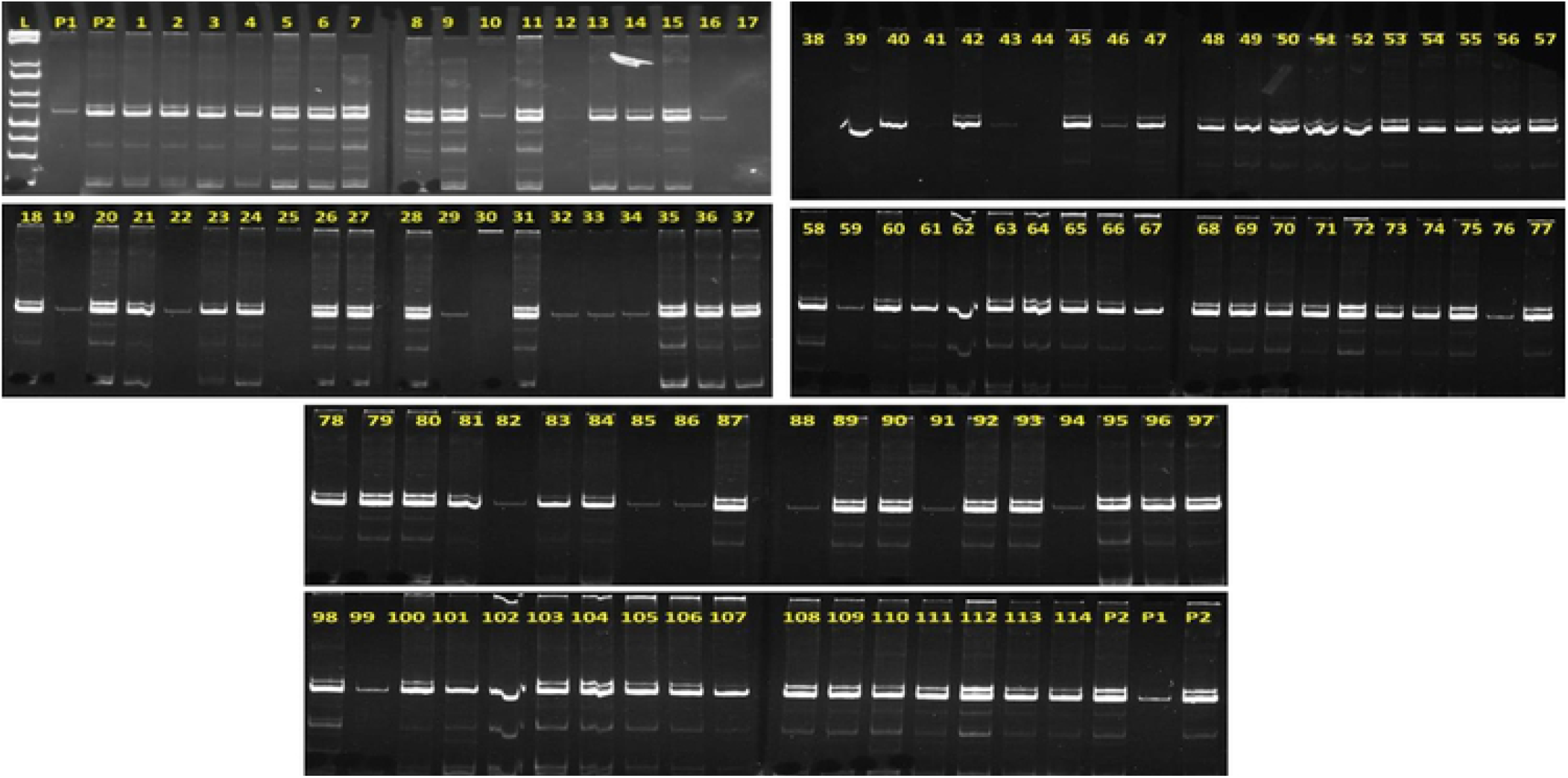
PAGE images showing validation of polymorphic SSR markers upon RILs.

## Conclusion

Next generation sequencing is a powerful tool for rapid collection of vast amount of RNASeq data from organisms and detailed analysis of the complete set of transcripts for understanding their developmental stages and physiological conditions [76–78]. The analysis workflow enables the discovery of unique and candidate genes having significant roles in the molecular and biological processes particularly for the non-model species like eggplant. Moreover, a large number of SSR and SNP markers can also be generated from the assembly [79, 80]. Since the genome of several species of eggplant still remains lagged behind the other vegetable crops like tomato and potato, RNASeq technique would bring this crop a stage up towards deciphering the molecular mechanism underlying defence, developmental and physiological responses through the structural, functional, and phylogenomics approach.

The present report is the first transcriptome sequencing assembly information of the cultivar Ramnagar Giant which is least characterized species from *melongenae* complex of eggplant. The cultivar can be useful for breeding purpose for development of high yielding varieties. The Ramnagar Giant x W-4 developed RILs population would be further utilized for QTL mapping of horticultural and domestication traits in eggplant. High morphological variation was observed among the RILs. Analysis of polymorphism and variation in this population would be useful for tracing the significant QTLs distributed along the eggplant genome. Moreover, a huge number of SSR markers developed through RNA*Seq* in the present study will be useful to provide a good platform for mapping and diversity studies on intra- and inter-specific population of eggplant genotypes. Furthermore, the markers will also be useful for elucidating genetic relationship among cultivated eggplant and their wild relatives. Lastly, the results assembled through the RNA*Seq* of the two species are enriched with high accuracy and reliable for future for identification of unique and novel candidate genes and development of more SSR and SNP markers with implications to modern marker-assisted breeding in eggplant. Overall, the comprehensive datasets obtained through the Ramnagar Giant and W-4 transcriptomes will be used for QTL mapping and serve as a reference for further analysis of disease resistance candidate genes and broaden the understanding of this agriculturally important vegetable crop.

## Acknowledgements

Indian Council of Agricultural Research (ICAR), New Delhi is acknowledged for funding this research. Authors are thankful to the Director, ICAR-IIVR, Varanasi, Head, Division of Crop Improvement, ICAR-IIVR, Varanasi and The Coordinator, School of Biotechnology, BHU, Varanasi for providing necessary facilities to carry out the present analysis.

## Author Contributions

Conceptualization of experiment: SKT, MS, PSN and BS Formal analysis: SKT, PM, SS, VKS. Experimental validation: PM and SPK. Contributed reagents/materials/analysis tools: SKT, VKS, MS, PSN and BS. Supervision: SKT. Original draft writing: PM, SPK and SKT. Review and editing of original draft: VKS, and KNT.

## Supporting Information

**S1 Table. Primary data statistics report.**

**S2 Table. *S. melongena* and *S. incanum* SSR details and primer sequences.**

**S3 Table. *S. melongena* and *S. incanum* SSR analysis data.**

**S4 Table. *S. melongena* and *S. incanum* SNV details.**

**S5 Table. *S. melongena* and *S. incanum* quantification data.**

**S6 Table. *S. melongena* transcripts annotation report.**

**S7 Table. *S. incanum* transcripts annotation report.**

**S8 Table. KEEG pathway analysis report.**

**S9 Table. Other analysed data.**

**S10 Table. *S. melongena* and *S. incanum* TFs summery.**

